# A novel widespread bacterial structure related to the flagellar type III secretion system

**DOI:** 10.1101/2021.09.03.458937

**Authors:** Mohammed Kaplan, Catherine M Oikonomou, Cecily R. Wood, Georges Chreifi, Debnath Ghosal, Megan J Dobro, Qing Yao, Alasdair McDowall, Ariane Briegel, Morgan Beeby, Yi-Wei Chang, Carrie L. Shaffer, Grant J. Jensen

## Abstract

The flagellar type III secretion system (fT3SS) is a suite of membrane-embedded and cytoplasmic proteins responsible for building the bacterial flagellar motility machinery. Homologous proteins form the injectisome machinery bacteria use to deliver effector proteins into eukaryotic cells, and other family members have recently been reported to be involved in the formation of membrane nanotubes. Here we describe a novel, ubiquitous and evolutionarily widespread hat-shaped structure embedded in the inner membrane of bacteria, of yet-unidentified function, that is related to the fT3SS, adding to the already rich repertoire of this family of nanomachines.

---

Type III secretion systems (T3SS) assemble bacterial machinery with diverse functions. In addition to forming the flagellum and the injectisome, they have also been reported recently to be involved in the formation of membrane tubes [1]. The flagellar type III secretion system (fT3SS) consists of a cytoplasmic part containing an ATPase and an inner-membrane (IM)-embedded part known as the core complex (fT3SScc). The fT3SScc consists of five proteins (FliP, FliQ, FliR, FlhB and FlhA), with another protein, FliO, required for assembly but which does not form part of the complex [2,3]. Initially, FliP forms a pentameric platform on which FliQ, FliR and FlhB assemble to create a FliP_5_FliQ_4_FliR_1_FlhB_1_ subcomplex upon which an FlhA ring is built [4].

While using electron cryo-tomography (cryo-ET) to study the process of flagellar assembly in *Helicobacter pylori*, we identified a periplasmic hat-shaped structure embedded in the inner membrane (IM) of the cell (Fig. 1). The structure was abundant and, in contrast to the polar flagella of this species, did not show any preferred spatial localization in the cell (e.g., not exclusively at the cell pole). Carefully reexamining tens of thousands of cryotomograms of other, phylogenetically-diverse bacterial species our lab has imaged over the past 15 years, we found that this hat-like structure is widespread in diverse Gram-negative and Gram-positive bacteria (Fig. 2; see also Supporting Figure S1). In many cases, we observed multiple hat-like structures (up to 10 in some cells) distributed around the cell (see Movie S1 for an example from an *E. coli* cell that was partially lysed, enhancing visibility of periplasmic structures). Subtomogram averages of the structure from different species revealed conserved characteristics: a hat-shaped part in the periplasm and two cytoplasmic densities beneath (Fig. 2). In general, the periplasmic hat-like portion had a diameter of ∼24-26 nm at its widest point at the outer surface of the IM. The cytoplasmic densities were absent in the averages from three species: *Pseudoalteromonas luteoviolacea, Hylemonella gracilis* and *Bacillus subtilis*. The absence of these densities in *P. luteoviolacea* and *B. subtilis* is likely due to the fact that these were lysed and not intact cells. We also observed that the cytoplasmic density did not resolve into two distinct sections in all species.

**Figure 1:**
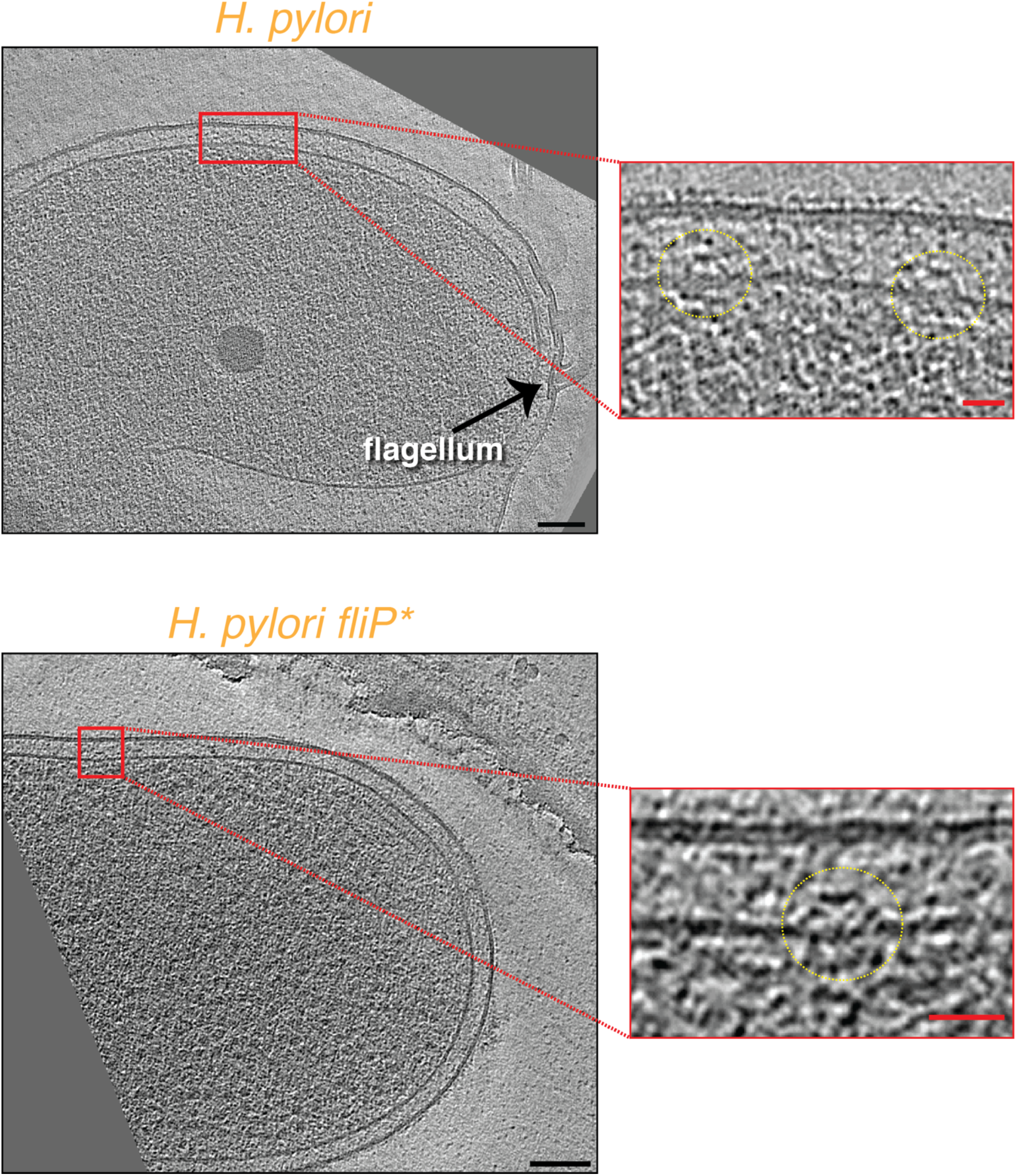
Identification of a novel hat-like complex in *H. pylori*. **A & B)** slices through electron cryotomograms of *H. pylori* (A) or *H. pylori fliP*\* (B) cells showing the presence of hat-like complexes (enlarged in red boxes). Black scale bars 100 nm, red scale bars 25 nm.

**Figure 2:**
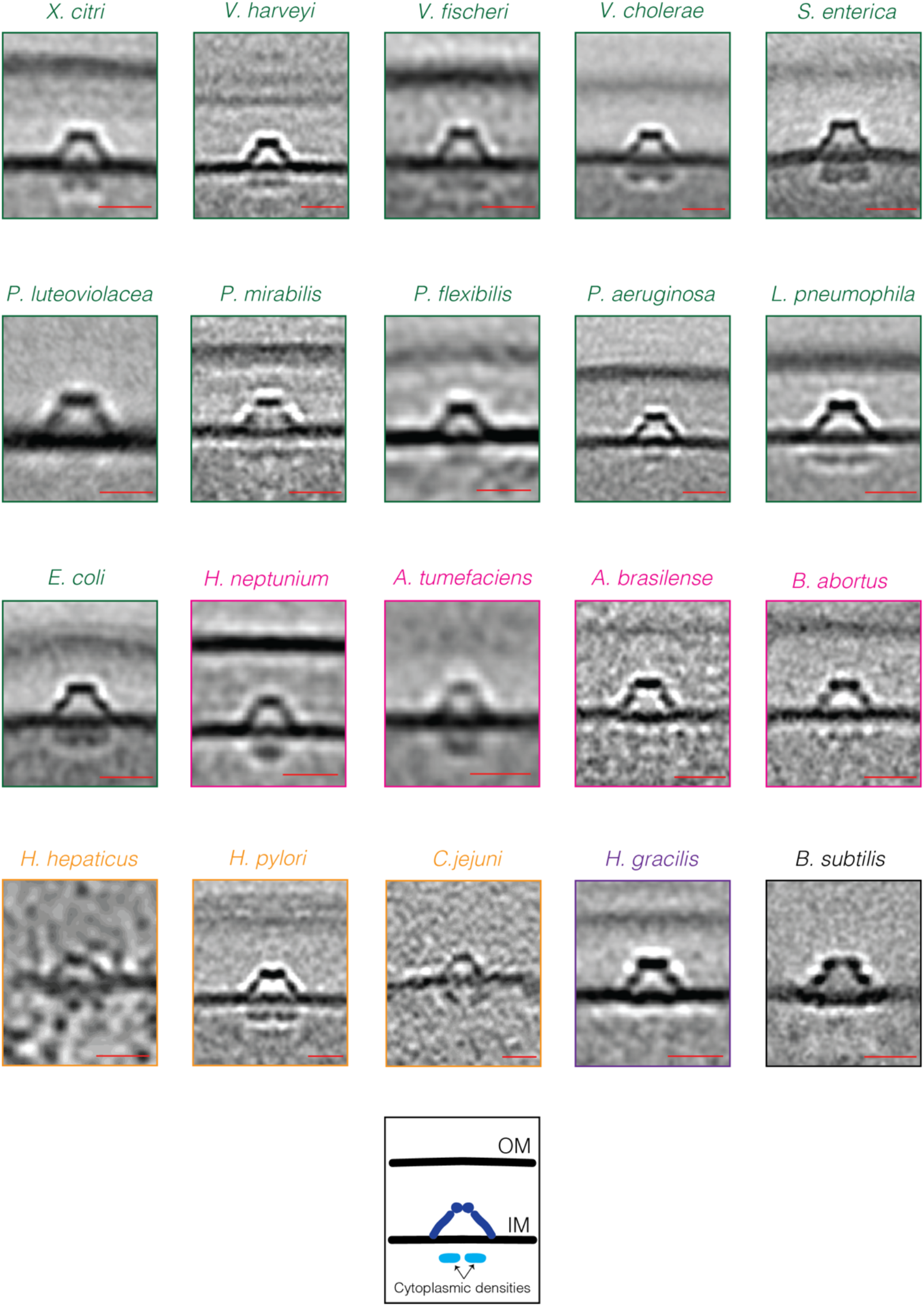
The hat-like complex is a widespread bacterial structure. A gallery of the hat-like complex in different bacterial species (*Xanthomonas citri, Vibrio harveyi, V. fischeri, V. cholerae, Salmonella enterica, Pseudoalteromonas luteoviolacea, Proteus mirabilis, Pseudomonas flexibilis, P. aeruginosa, Legionella pneumophila, Escherichia coli, Hyphomonas neptunium, Agrobacterium tumefaciens, Azospirillum brasilense, Brucella abortus, Helicobacter hepaticus, H. pylori, Campylobacter jejuni, Hylemonella gracilis* and *Bacillus subtilis*). Sub-tomogram averages are shown, except for *C. jejuni* and *H. hepaticus*, where not enough data was available for averaging so single tomographic slices are shown. Color coding indicates taxonomic class: green, Gammaproteobacteria; pink, Alphaproteobacteria; yellow, Epsilonproteobacteria; purple, Betaproteobacteria; and black, Bacilli. Scale bars are 20 nm.

We also identified the same structure in several *H. pylori* flagellar mutants: *fliP*\*, Δ*flgS fliP*\*, Δ*fliM fliP*\*, Δ*fliG fliP*\*, Δ*fliO fliP*\*, and Δ*fliQ fliP*\*. The *H. pylori fliP*\*strain contains a naturally-occurring point mutation that disrupts the function of FliP [5] and prevents the assembly of the fT3SScc (manuscript in preparation). The other mutants remove additional fT3SScc proteins (Δ*fliO* and Δ*fliQ*), flagellar basal body proteins (Δ*fliM* and Δ*fliG*), or the tyrosine kinase responsible for expression of the class II flagellar genes (Δ*flgS*) [6]. Curiously, in all of these mutants the diameter of the hat-like density was reduced to only ∼20 nm (at its widest part) and the two cytoplasmic densities were missing (or less well resolved) (Fig. 3 A-G). This difference was not due to decreased resolution, since more particles were averaged than from wild-type cells (see Materials and Methods). This observation suggested to us that the hat-like structure is related to the fT3SScc. Indeed the general shape is reminiscent of the MS-ring of the flagellar motor, and we observed the disappearance of two similar cytoplasmic densities in the motor (corresponding to FlhA_C_) in the same mutants while studying flagellar assembly (manuscript in preparation). The reduced width of the hat-like structure in *fliP*\* cells is also reminiscent of the reduced width of flagellar complexes in the absence of the fT3SScc [7].

**Figure 3:**
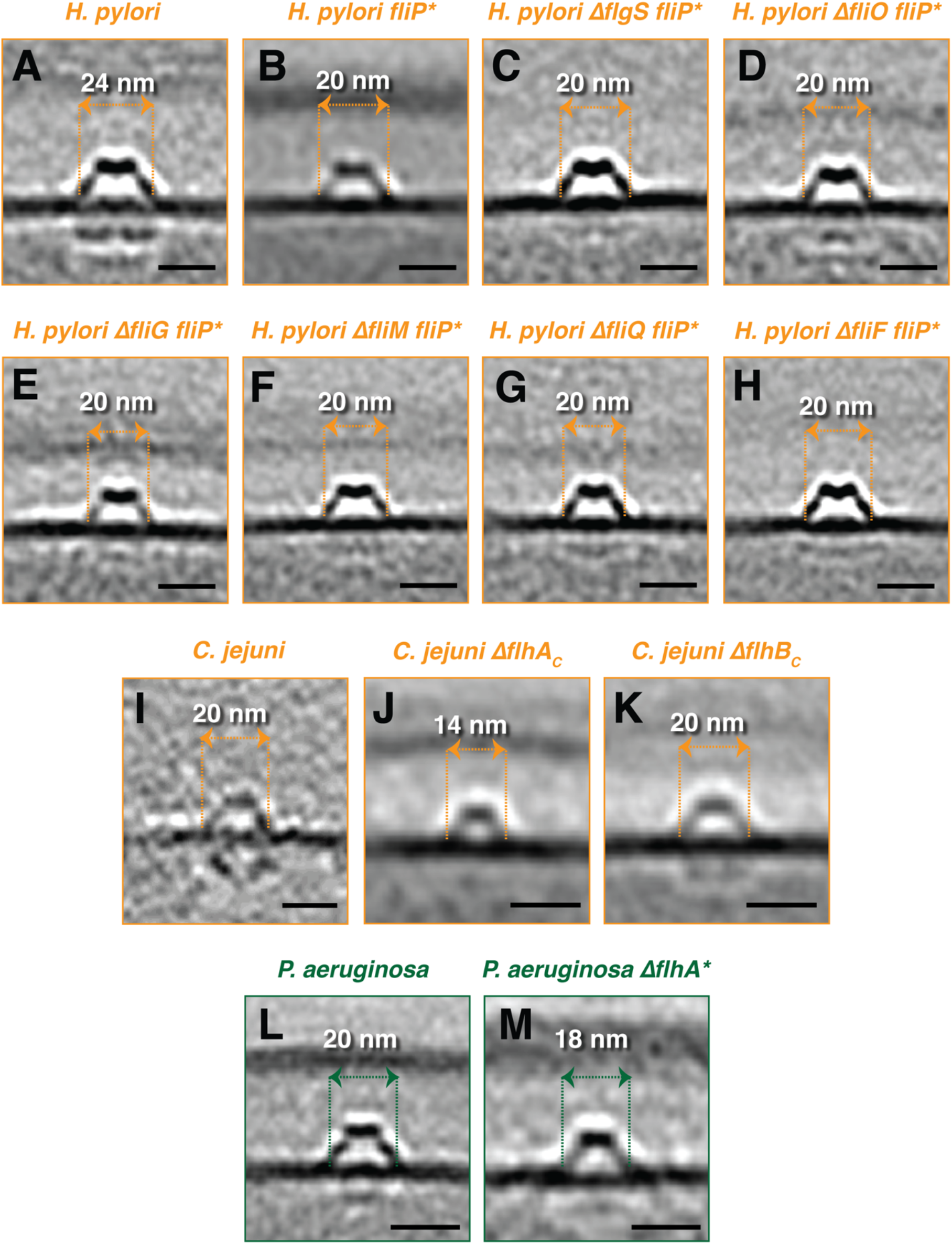
The effect of various flagellar-related mutations on the hat-like complex. Central slices through sub-tomogram averages (except (I), where a single tomographic slice is shown) of the hat-like complex in wild-type cells and the indicated mutants of *H. pylori* **(A-H)**, *C. jejuni* **(I-K)** and *P. aeruginosa* **(L & M)**. Scale bars are 20 nm.

To explore this relationship, we examined *Campylobacter jejuni* mutants of other fTSScc proteins. These included mutants of the C-terminal domains of FlhA (Δ*flhAc*) and FlhB (Δ*flhBc*) [8]. In Δ*flhAc* cells, compared to wild-type, the periplasmic hat-like part was again smaller in diameter and the two cytoplasmic densities disappeared (Fig. 3 I & J). In contrast, the hat-like structure in Δ*flhBc* cells was indistinguishable from the wild-type complex both in diameter and the presence of the associated cytoplasmic densities (Fig. 3 K). This is not too surprising, since, unlike the large pentameric FliP ring or the nonameric FlhA ring, FlhB is a small protein present in a monomeric form in the fT3SScc. Although the absence of the C-terminus of FlhB renders the fT3SS non-functional (no full flagella assemble in Δ*flhBc C. jejuni* [8]), the fT3SScc can still assemble (manuscript in preparation). To confirm the generality of the relationship between the fT3SScc and the hat-like complex, we imaged an *flhA* mutant in *P. aeruginosa* (*flhA*\*, obtained from a transposon insertion mutant library). Here also, the hat-like structure was smaller in size and lacked clear cytoplasmic densities compared to wild-type (Fig. 3 L & M).

Based on the apparent relationship between the fT3SScc and the hat-like structure, we hypothesized that the novel complex is formed by the flagellar MS-ring protein, FliF, adopting a different, more closed conformation than that seen in the fully-assembled flagellar motor. Hence, we generated and imaged a Δ*fliF* mutant in the *H. pylori fliP*\* background. However, the hat-like complex was still present in this mutant, indicating that it is not formed by FliF (Fig. 3 H). Thus our observations suggest that while the cytoplasmic densities of the complex could be FlhA_C_, the periplasmic density is not formed by FliF or any of the fT3SScc proteins. Of course, it is also possible that FlhA_C_ does not directly constitute the cytoplasmic densities, but rather that the fT3SScc proteins are regulating the expression (or localization) of another protein(s) that does.

One possibility is that the hat-like structure we discovered here might represent an as-yet unidentified scaffold that helps the fT3SScc assemble. If this structure is some sort of scaffold and the fT3SScc (or part of it) assembles within the hat-like portion and if the cytoplasmic densities are FlhA_C_, this would explain the disappearance of the cytoplasmic densities and the smaller diameter of the periplasmic portion in *fliP*\* and *flhA* mutants. The second possibility is that fT3SScc proteins in some way regulate other proteins which themselves form the hat-like complex. Such a regulatory role has been indicated previously for one of the fT3SScc proteins, FliO, which is responsible for the optimal expression of other flagellar genes [9].

Whatever the function of this hat-like complex, it joins the already-rich repertoire of the (f)T3SS, which has roles in flagellar motility, protein translocation and possibly membrane nanotube formation. Whether the hat-like structure is connected to any of these functions or plays another, yet-unidentified role remains to be elucidated. It is also possible that the apparently ancient structure may have diverged to serve different functions in different species.

## Supporting Information for

### Supporting Figures

**Figure S1:**
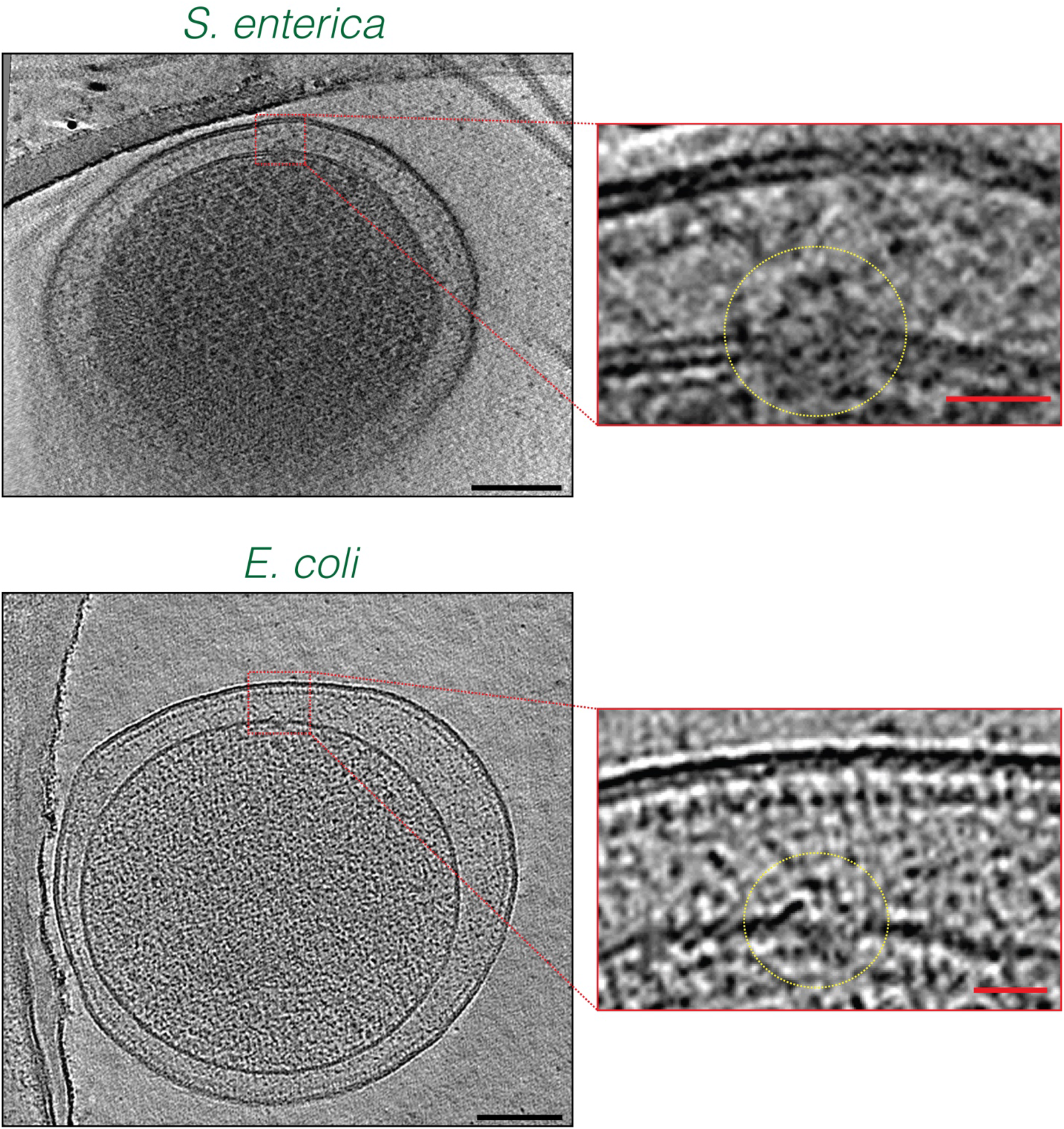
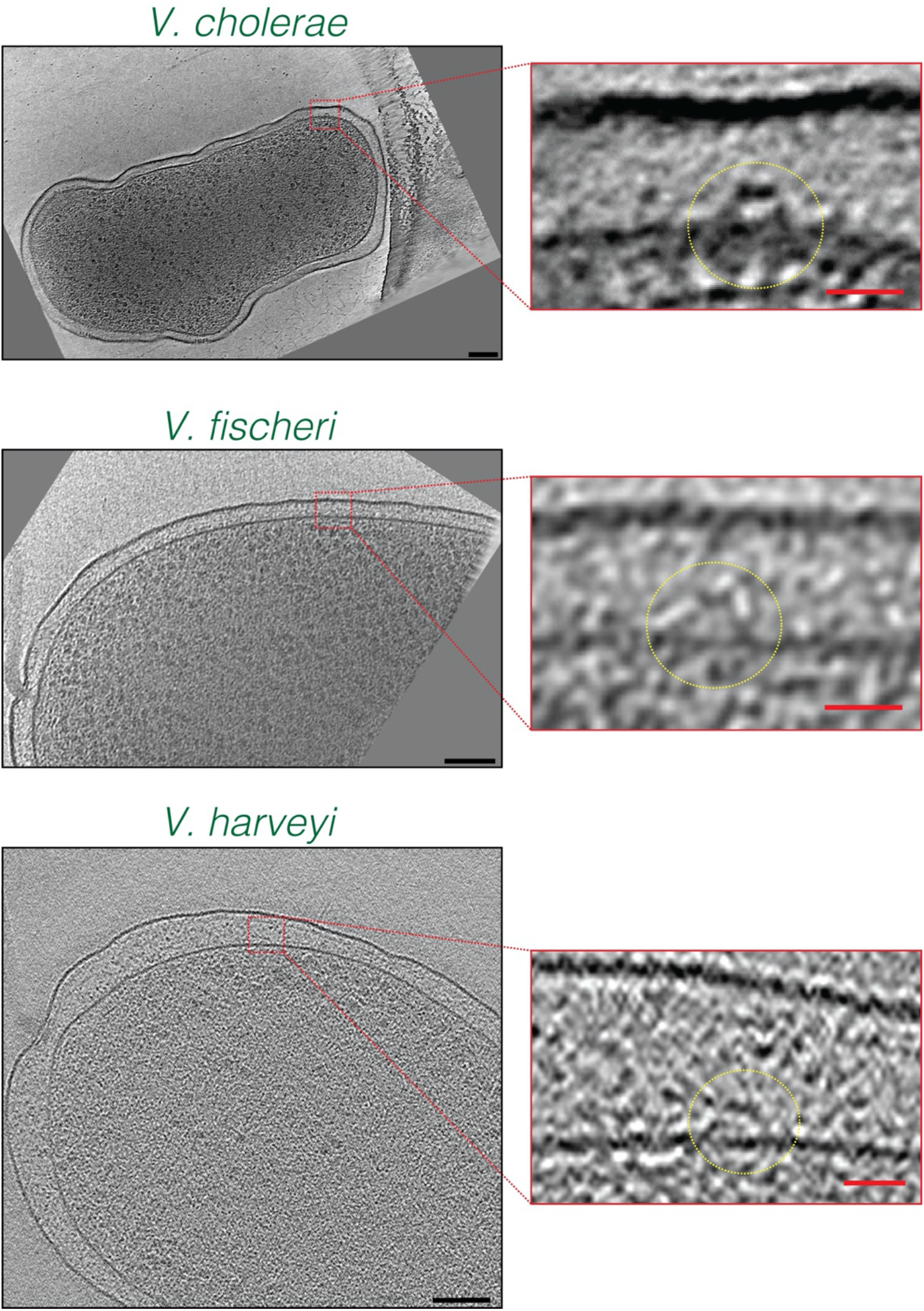
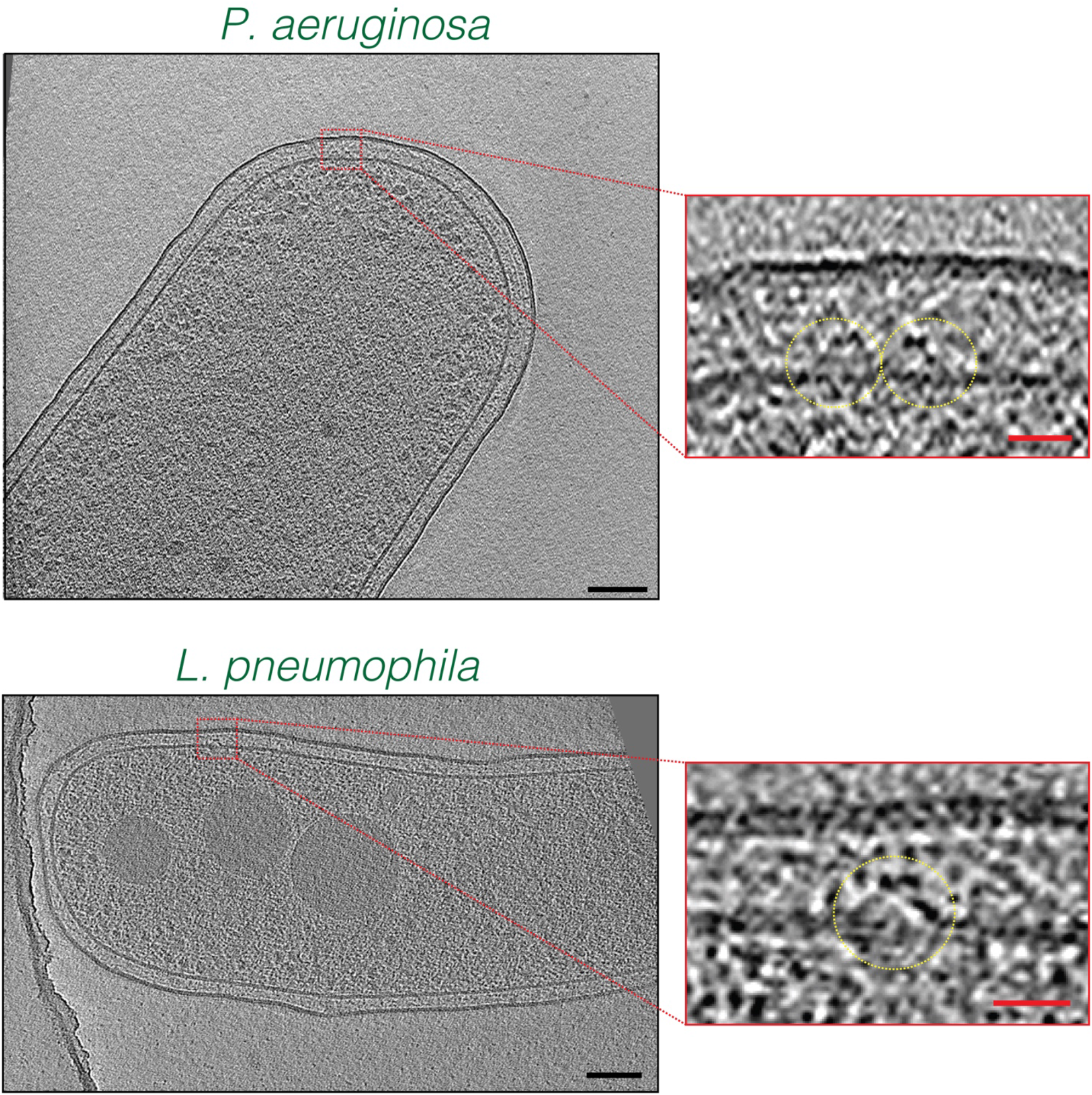
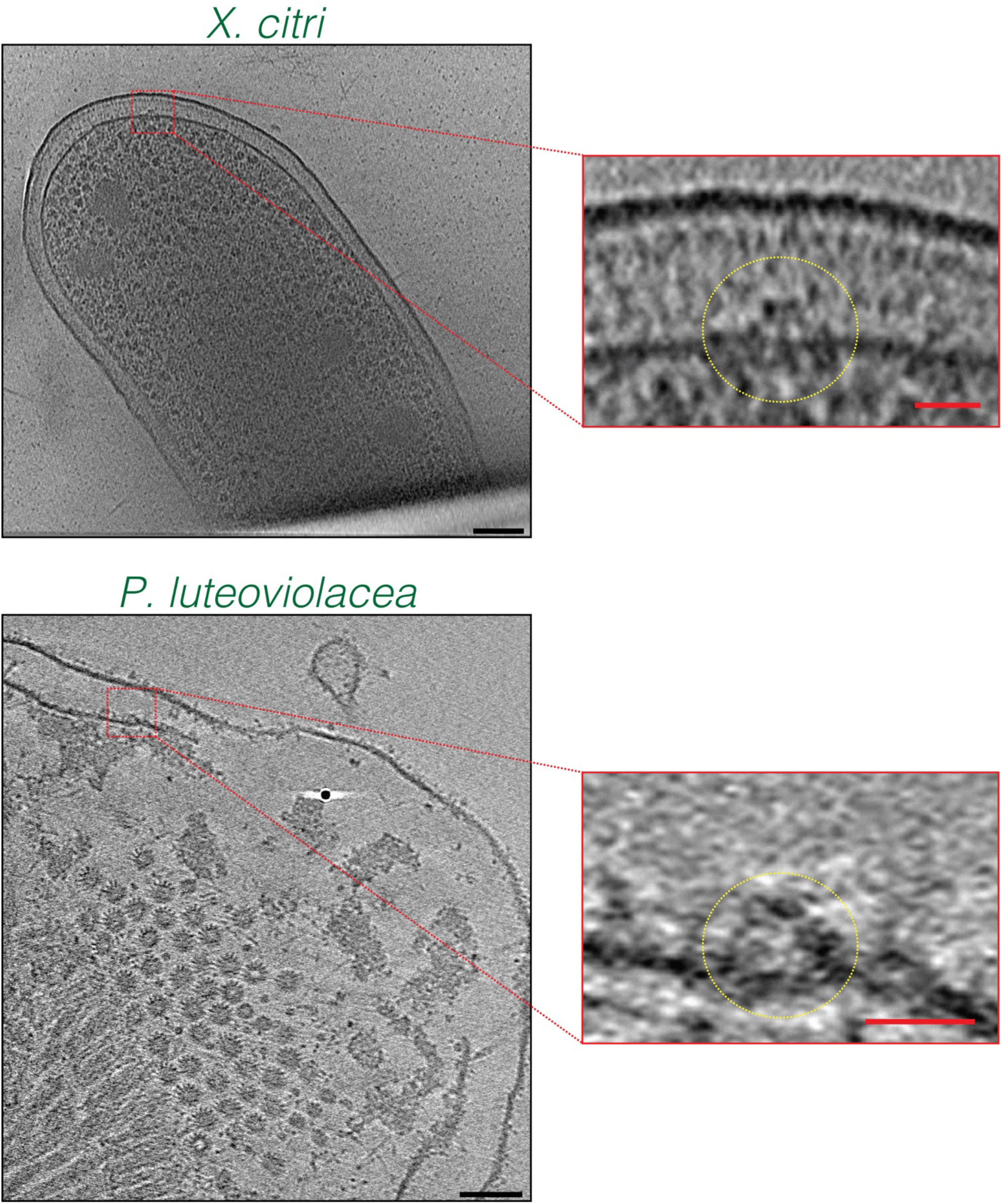
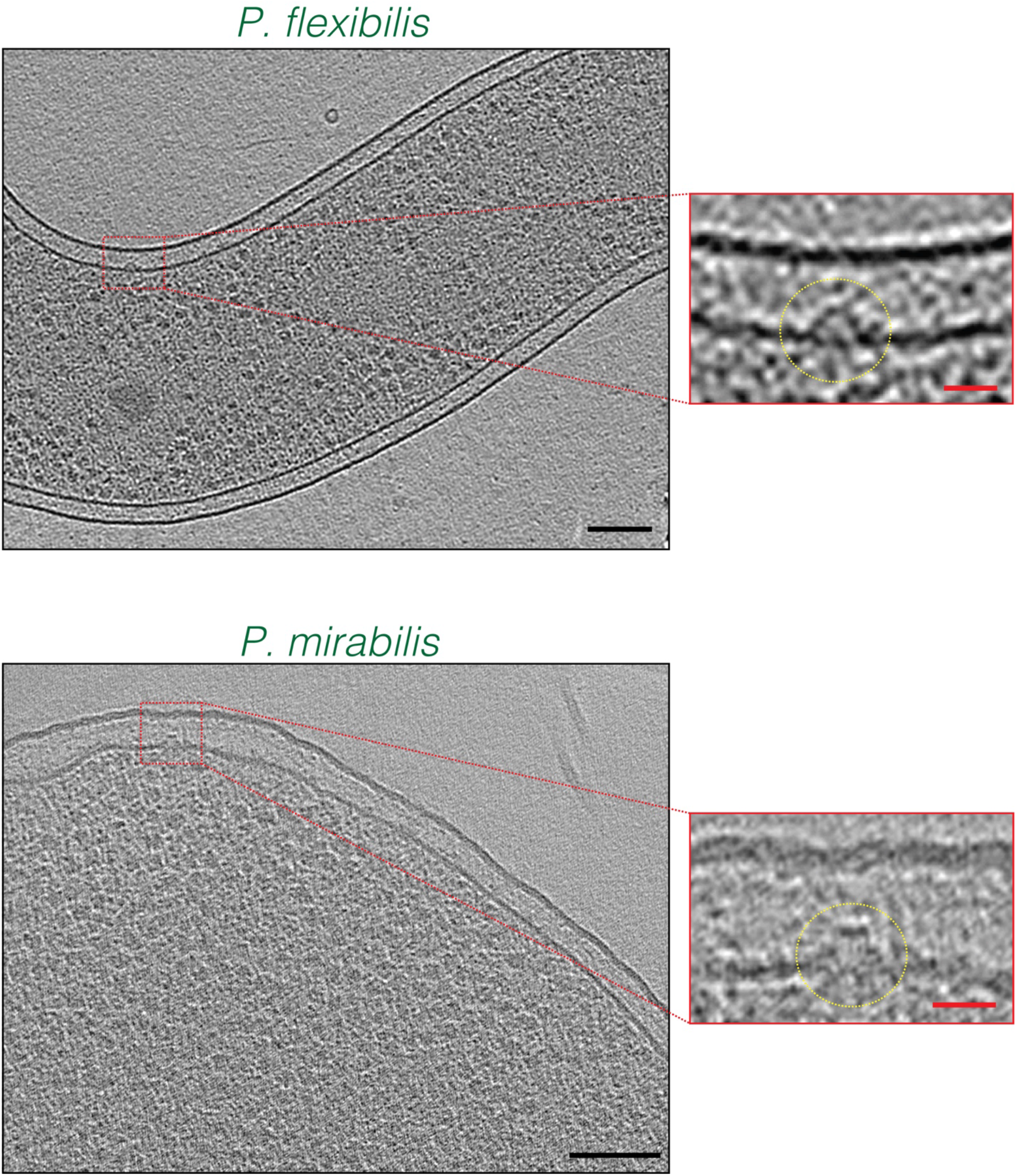
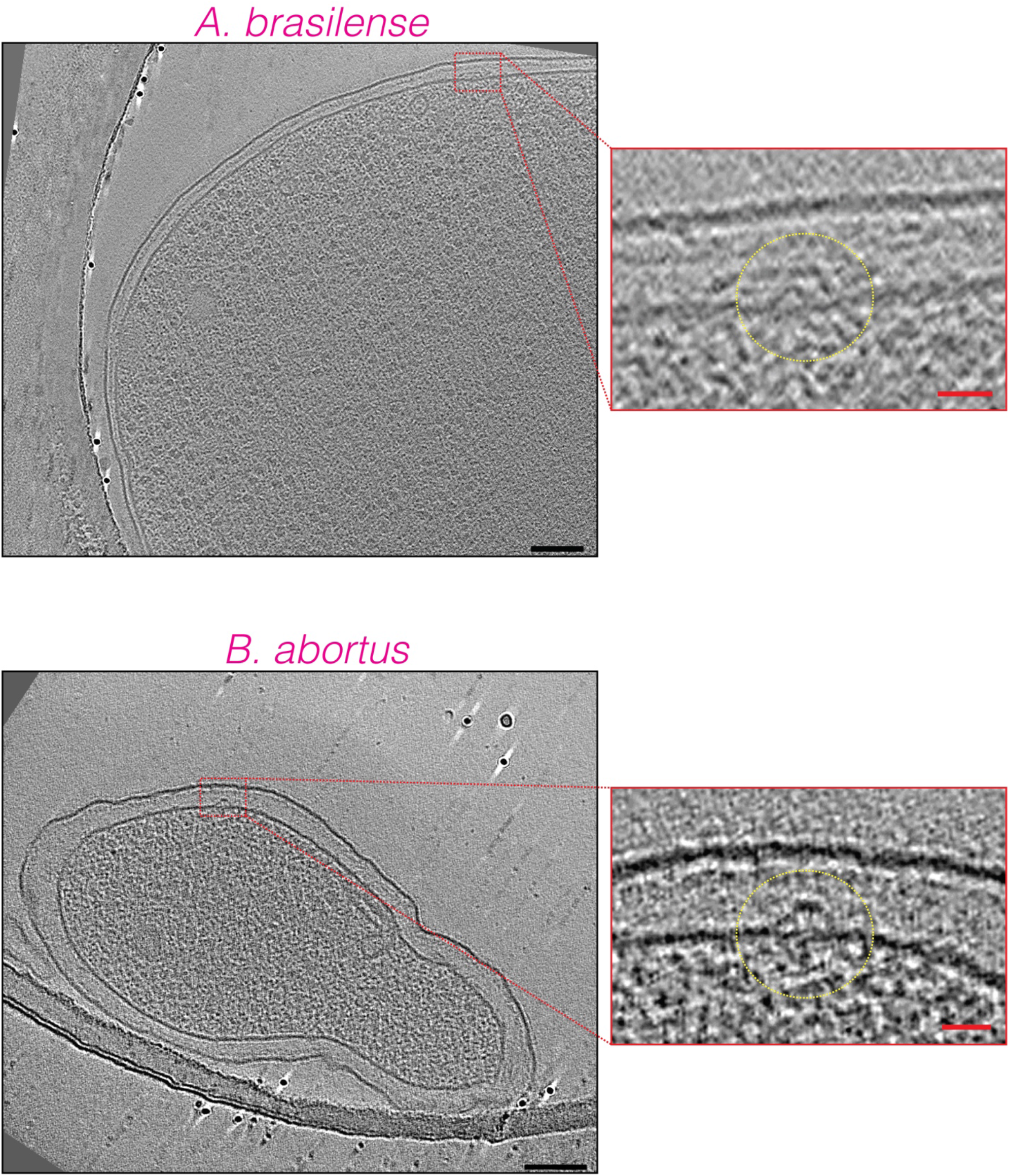
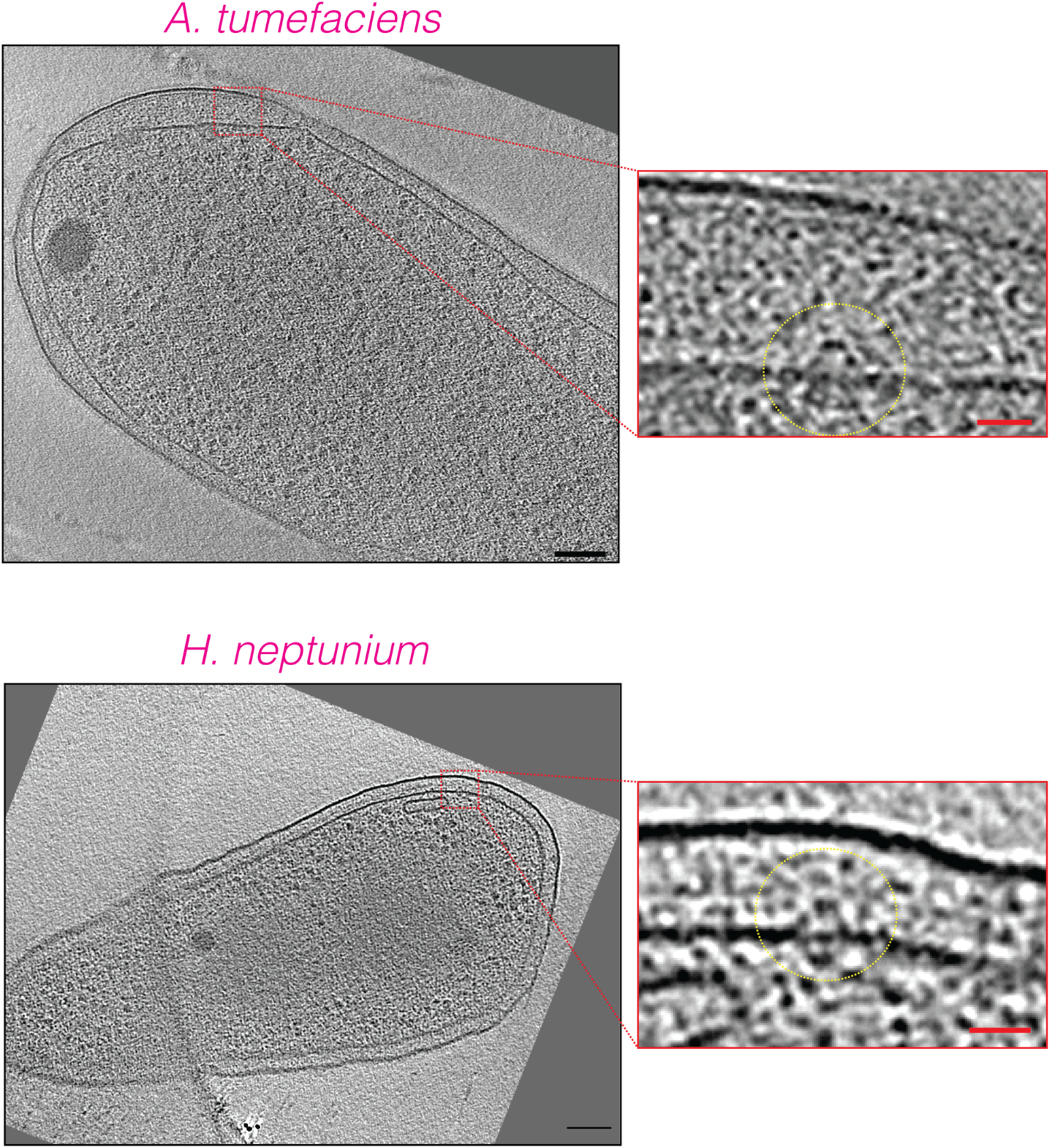
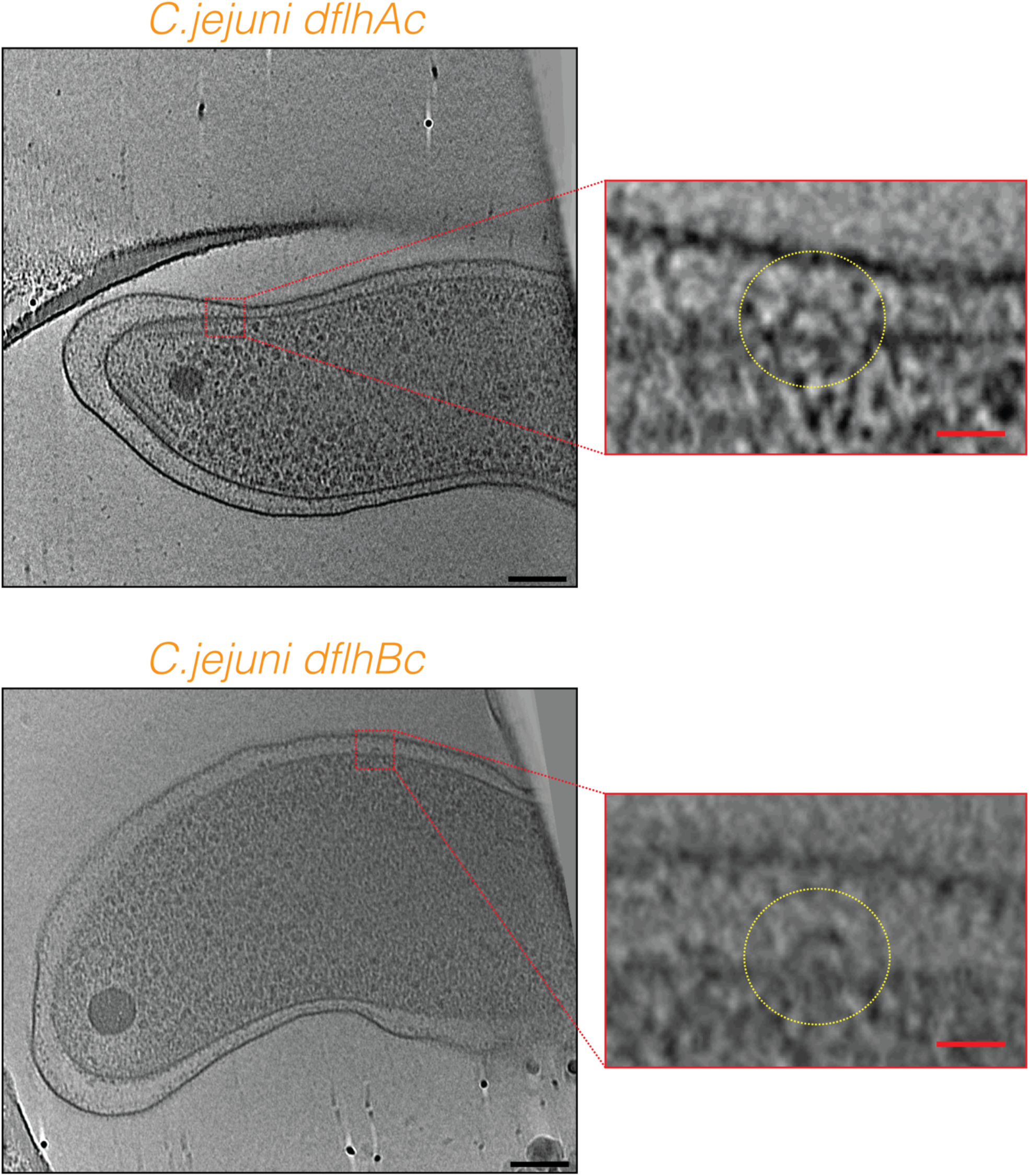
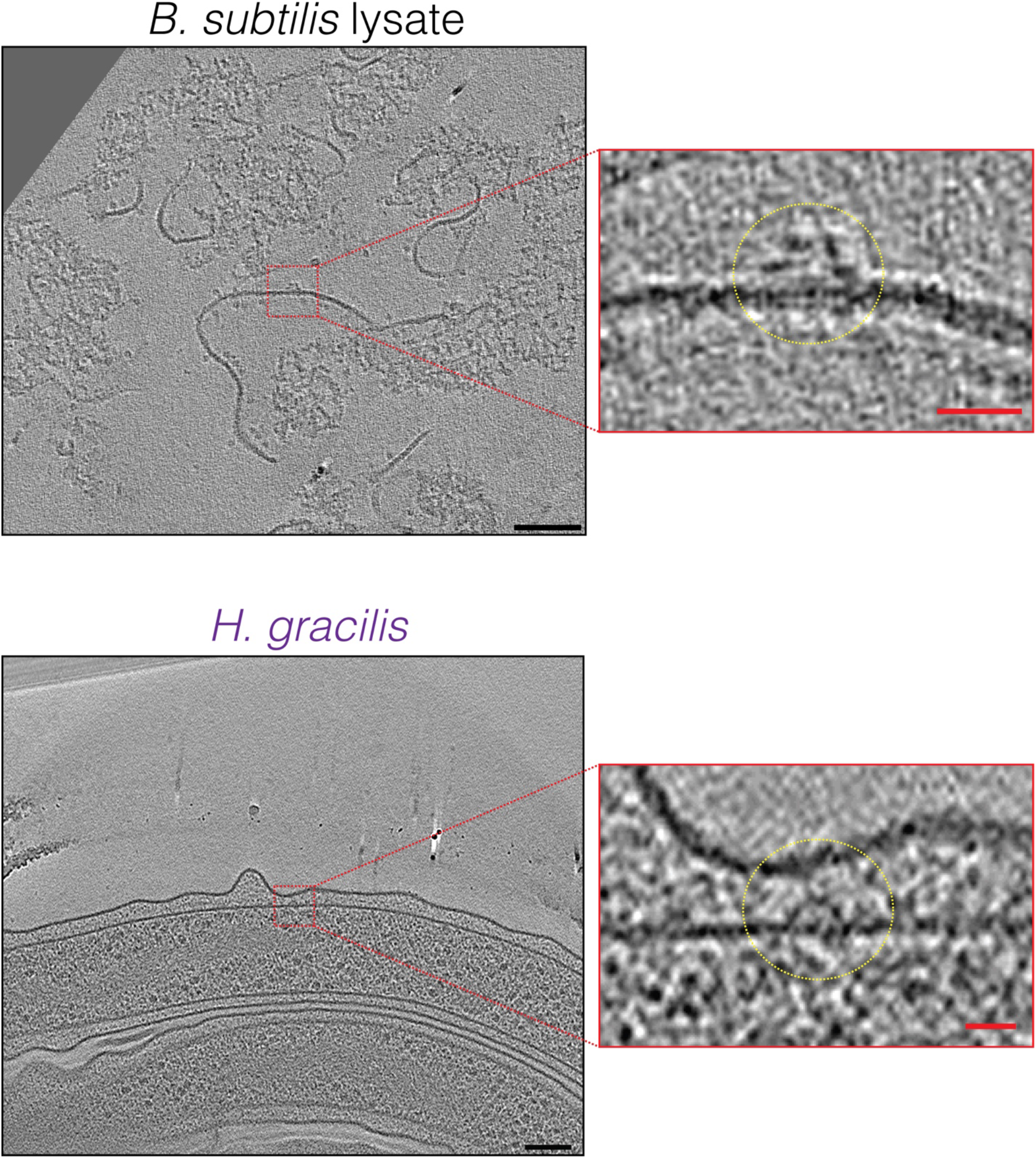
Slices through electron cryotomograms of various bacterial species highlighting the presence of hat-like complexes (dotted yellow circles in the enlarged views). Black scale bars 100 nm, red scale bars 20 nm.

### Movie S1

An electron cryotomogram of a partially-lysed *E. coli* cell highlighting the presence of multiple hat-like complexes in the inner membrane (indicated by red circles).

### Extended Materials and Methods

#### Strains and growth conditions

*E. coli* cells were grown as described in ref. [1]. *X. citri* cells were grown in 2xTY medium for 14 hours to stationary phase. *V. cholerae, V. harveyi* and *V. fischeri* were grown as previously described [2]. *P. luteoviolacea* were grown as described in ref. [3]. *P. mirabilis* were grown as described in ref. [4]. *P. aeruginosa* were grown in LB medium at 37° C overnight. The *P. aeruginosa flhA*\* mutant was obtained from a transposon library (mutant number 3296 from the non-redundant library http://pa14.mgh.harvard.edu/cgi-bin/pa14/downloads.cgi) from Dianne Newman’s lab at Caltech. *L. pneumophila* were grown as described in ref. [5]. *S. enterica* were grown as in ref. [6]. *P. flexibilis* were grown in Lactose growth medium. *H. neptunium* were grown to exponential phase in PYE medium. *A. tumefaciens* wild-type cells with plasmid-borne VirC1-GFP translational fusion under control of the VirB promoter were grown in AB medium with 150-300 ug/ml of kanamycin. *A. brasilense* and *B. abortus* were grown as described in ref. [7]. *H. hepaticus* ATCC 51449 and *H gracilis* were grown as described in ref. [1,8] *C. jejuni* and its mutants were grown as described in ref. [6,9,10]. *B. subtilis* protoplasts were prepared using lysozyme using a protocol based on ref. [11]. A motile revertant *H. pylori* 26695 isolate was selected by serial passage in Brucella broth supplemented with 10% heat inactivated fetal bovine serum at 37° C, 5% CO_2_ for 4 days until cultures reached an OD_600_ ∼ 0.4. Non-motile *H. pylori fliP*\* mutants were propagated on TSAII blood agar plates (BD Biosciences) at 37 °C, 5% CO_2_ for either 24 or 48 h prior to collection with a sterile cotton swab for grid preparation. *Helicobacter pylori* mutants (Δ*fliM fliP*\*, Δ*fliO fliP*\*, Δ*flgS fliP*\*, Δ*fliG fliP*\*, Δ*fliQ fliP*\*) were grown directly from glycerol stocks on sheep blood agar at 37 °C with 5% CO_2_ for 48 hours. Then, the cells were either collected from the plate using a cotton swab and dissolved in PBS and spun down and plunge-frozen directly, or the cells were spread on a new plate and allowed to grow for 24 hours under the same conditions before plunge-freezing. No difference could be discerned between the two samples by cryo-ET.

#### *H. pylori* mutagenesis

Flagellar mutants were generated in the non-motile *H. pylori* 26695 background as previously described [12]. Briefly, constructs were generated to replace the coding region of the gene of interest with an in-frame, non-polar kanamycin resistance cassette. The target gene and approximately 500 base pairs (bp) upstream and downstream of flanking regions were amplified and cloned into pGEM T-Easy (Promega). This construct was used as a template for inverse PCR to remove the majority of the target gene coding region and to introduce incompatible restriction sites for directional cloning. A kanamycin resistance cassette driven by a promoter transcribed in the same direction as the endogenous operon was cloned into the ligated inverse PCR plasmid. *H. pylori* 26695 was transformed via natural competence, and single colonies resistant to kanamycin (12.5 µg/ml) were selected. PCR was used to verify that the kanamycin resistance cassette had inserted into the target locus in the same orientation as operon transcription.

#### Electron cryo-tomography sample preparation and imaging

Sample preparation for cryo-ET imaging was done as described in references [2,13,14]. Total cumulative electron dose used for each tilt-series in each species was:

| Species name | Class | Cumulative electron dose (e<sup>-</sup>/ Å<sup>2</sup>) |
| --- | --- | --- |
| *Xanthomonas citri* | Gammaproteobacteria | 120 |
| *Vibrio harveyi* | Gammaproteobacteria | 160 |
| *Vibrio fischeri* | Gammaproteobacteria | 150 |
| *Vibrio cholerae* | Gammaproteobacteria | 160 |
| *Salmonella enterica* | Gammaproteobacteria | 200 |
| *Pseudoalteromonas*<br>*luteoviolacea* | Gammaproteobacteria | 180 |
| *Proteus mirabilis* | Gammaproteobacteria | 160 |
| *Pseudomonas flexibilis* | Gammaproteobacteria | 100 |
| *Pseudomonas fluorescens* | Gammaproteobacteria | 200 |
| *Pseudomonas aeruginosa* | Gammaproteobacteria | 170 |
| *Legionella pneumophila* | Gammaproteobacteria | 100 |
| *Escherichia coli* | Gammaproteobacteria | 130 |
| *Shewanella oneidensis* MR1 | Gammaproteobacteria | 150 |
| *Hyphomonas neptunium* | Alphaproteobacteria | 180 |
| *Agrobacterium tumefaciens* | Alphaproteobacteria | 200 |
| *Azospirillum brasilense* | Alphaproteobacteria | 200 |
| *Brucella abortus* | Alphaproteobacteria | 160 |
| *Helicobacter hepaticus* | Epsilonproteobacteria | 200 |
| *Helicobacter pylori* | Epsilonproteobacteria | 120-130 |
| *Campylobacter jejuni* | Epsilonproteobacteria | 200 |
| *Hylemonella gracilis* | Betaproteobacteria | 75 |
| *Bacillus subtilis* | Bacilli | 160 |

#### Image processing and subtomogram averaging

Three-dimensional reconstructions of tilt-series were performed either automatically through the RAPTOR pipeline used in the Jensen lab [15] or with the IMOD software package [16]. Subtomogram averaging was done using the PEET program [17] with a 2-fold symmetrization applied along the particle Y-axis. The number of particles that were averaged are:

| Figure | Number of particles |
| --- | --- |
| Figure 2 *X. citri* | 21 |
| Figure 2 *V. harveyi* | 46 |
| Figure 2 *V. fischeri* | 19 |
| Figure 2 *V. cholerae* | 58 |
| Figure 2 *S. enterica* | 38 |
| Figure 2 *P. luteoviolacea* | 50 |
| Figure 2 *P. mirabilis* | 18 |
| Figure 2 *P. flexibilis* | 23 |
| Figure 2 *P. aeruginosa* | 78 |
| Figure 2 *L. pneumophila* | 149 |
| Figure 2 *E. coli* | 31 |
| Figure 2 *H. neptunium* | 23 |
| Figure 2 *A. tumefaciens* | 29 |
| Figure 2 *A. brasilense* | 20 |
| Figure 2 *B. abortus* | 41 |
| Figure 2 *H. pylori* | 26 |
| Figure 2 *H. gracilis* | 30 |
| Figure 2 *B. subtilis* | 30 |
| Figure 3A | 26 |
| Figure 3B | 118 |
| Figure 3C | 60 |
| Figure 3D | 29 |
| Figure 3E | 27 |
| Figure 3F | 50 |
| Figure 3G | 42 |
| Figure 3H | 146 |
| Figure 3J | 36 |
| Figure 3K | 37 |
| Figure 3L | 78 |
| Figure 3M | 17 |

## Acknowledgements

This project was funded by the NIH (grant R01 AI127401 to G.J.J., and P20 GM130456 and P30 GM110787 to C.L.S.) and a Baxter postdoctoral fellowship from Caltech to M.K. Cryo-ET work was done in the Beckman Institute Resource Center for Transmission Electron Microscopy at the California Institute of Technology. We are grateful to Prof. Marc Erhardt (Humboldt-Universität zu Berlin) for critically reading an initial version of this work. We thank Prof. Elitza I. Tocheva for collecting the *A. tumefaciens* data, Dr. Jian Shi for collecting the *H. neptunium* data, and Prof. Martin Pilhofer for collecting the *P. luteoviolacea* data.

## Notes

### Competing Interest Statement

The authors have declared no competing interest.

